# Shugoshin is essential for meiotic prophase checkpoints in *C. elegans*

**DOI:** 10.1101/258830

**Authors:** Tisha Bohr, Christian R. Nelson, Stefani Giacopazzi, Piero Lamelza, Needhi Bhalla

**Affiliations:** Department of Molecular, Cell and Developmental Biology, University of California, Santa Cruz, Santa Cruz, CA 95064

**Keywords:** meiosis, checkpoint, synapsis, DNA damage response, recombination, homologous chromosome, chromosomes axis, HORMADs

## Abstract

The conserved factor Shugoshin is dispensable in *C. elegans* for the two-step loss of sister chromatid cohesion that directs the proper segregation of meiotic chromosomes. We show that the *C. elegans* ortholog of Shugoshin, SGO-1, is required for checkpoint activity in meiotic prophase. This role in checkpoint function is similar to that of the meiotic chromosomal protein, HTP-3. Null *sgo-1* mutants exhibit additional phenotypes similar to that of a partial loss of function allele of HTP-3: premature synaptonemal complex disassembly, the activation of alternate DNA repair pathways and an inability to recruit a conserved effector of the DNA damage pathway, HUS-1. SGO-1 localizes to pre-meiotic nuclei, when HTP-3 is present but not yet loaded onto chromosome axes, suggesting an early role in regulating meiotic chromosome metabolism. We propose that SGO-1 acts during pre-meiotic replication to ensure fully functional meiotic chromosome architecture, rendering these chromosomes competent for checkpoint activity and normal progression of meiotic recombination. Given that most research on Shugoshin has been focused on its regulation of sister chromatid cohesion in meiosis, this novel role may be conserved but previously uncharacterized in other organisms. Further, our findings expand the repertoire of Shugoshin’s functions beyond coordinating regulatory activities at the centromere.

## Introduction

Sexually reproducing organisms rely on the specialized cell division, meiosis, to generate haploid gametes, such as sperm and eggs, so that diploidy is restored upon fertilization. To promote proper disjunction of meiotic chromosomes, homologs undergo a series of progressively intimate interactions during meiotic prophase. Chromosomes identify their unique homolog, pair, and stabilize pairing via the assembly of the synaptonemal complex (SC) in a process called synapsis. Interhomolog crossover recombination occurs in the context of synapsis to produce linkages, or chiasmata, that direct meiotic chromosome segregation (reviewed in [1]). Defects in pairing, synapsis or recombination can produce errors in meiotic chromosome segregation and gametes with too few or too many chromosomes, also referred to as aneuploidy. Fertilization of these defective gametes generates aneuploid embryos, which are often inviable. It is estimated that ~30% of miscarriages are the result of aneuploidy [2] and many developmental disorders, such as Down or Klinefelter’s syndromes, are the product of aneuploidy.

Meiotic chromosomes are structured by a variety of proteins so that they are competent for pairing, synapsis and interhomolog recombination. These include the cohesin complex, which mediates sister chromatid cohesion, and axis component proteins that assemble the linear axes of the SC (reviewed in [3]). In addition, cohesin and axis components are involved in meiotic prophase checkpoints that respond to errors by either stalling meiotic prophase progression or activating apoptosis to remove defective meiocytes [4–10]. A subset of these proteins, identified by a conserved domain called the HORMA domain, adopt structures reminiscent of the spindle checkpoint effector, Mad2, suggesting that meiotic HORMA domain containing proteins (HORMADs) might also control meiotic checkpoint signaling through the adoption of multiple conformations [11, 12]. In budding yeast there is a single meiotic HORMAD (Hop1) [13], in mice there are two (HORMAD1 and 2) [14] and in *C. elegans* there are four (HTP-3, HIM-3, HTP-1 and HTP-2) [15–18]. Why this family has expanded so dramatically in *C. elegans* is unknown.

To halve the chromosome complement, meiosis is composed of two rounds of chromosome segregation: meiosis I, in which homologous chromosomes segregate, and meiosis II, in which sister chromatids segregate. This segregation scheme necessitates a two-step loss of sister chromatid cohesion. Cohesin is removed distal to chiasmata to allow homologs to segregate during meiosis I while being partially maintained to enable sister chromatids to partition correctly during meiosis II. In organisms with point centromeres, this sequential loss of cohesion is regulated by Shugoshin [19]. Shugoshin protects cohesin at the centromere until meiosis II by recruiting the conserved phosphatase, PP2A, to antagonize the phosphorylation and removal of the cohesin complex [20, 21]. Some organisms, such as *C. elegans*, do not have a localized centromere. In this model organism, the two-step loss of cohesin is accomplished through an alternate regulatory mechanism that involves the ordered, asymmetric disassembly of SC components around a single crossover site in late meiotic prophase [22–24]. Further, attempts to attribute a meiotic role to the worm ortholog of Shugoshin, SGO-1, have been unsuccessful [22].

We report here that SGO-1 is essential for checkpoint function in meiotic prophase in *C. elegans*. A hypomorphic mutant allele of *sgo-1* abrogates the synapsis checkpoint that monitors whether homologous chromosomes have synapsed, while a null mutation abrogates both the synapsis checkpoint and the DNA damage response (DDR). However, unlike other characterized synapsis checkpoint components, SGO-1 does not inhibit synapsis, indicating it acts in an alternate pathway. Instead, SC disassembly is accelerated in null *sgo-1* mutants *(sgo-1[0]))*. Both this phenotype and the requirement for SGO-1 in both meiotic checkpoints are reminiscent of phenotypes displayed by a partial loss of function mutation in the conserved chromosome axis component and meiotic HORMA domain containing protein, HTP-3. Indeed, similar to HTP-3, SGO-1 is also required for preventing the activation of alternate DNA repair pathways during meiosis and the ability to recruit conserved effectors of the DDR, such as HUS-1, suggesting a role in meiotic axis morphogenesis. Consistent with an early role in this process, SGO-1 localizes to pre-meiotic nuclei that express HTP-3 but have not yet assembled chromosome axes. We propose that SGO-1 contributes to checkpoint functionality and the proper progression of meiotic recombination through its regulation of meiotic chromosome architecture, potentially through its role as a regulator of cohesins, and suggest that this role may be conserved.

## Results

### SGO-1 is required for both the synapsis checkpoint and the DNA damage response

Previous experiments showed that a hypomorphic allele of *sgo-1, sgo-1(tm2443)*, resulted in low levels of chromosome segregation defects during meiosis [22], suggesting that SGO-1 might play a role in meiotic checkpoint function. The *sgo-1(tm2443)* allele results in a frameshift at amino acid 157 to generate a stop codon after 30 out-of-frame codons (Figure 1A) [25]. This results in a truncated protein product that is expressed at levels similar to that of full-length SGO-1 in wildtype animals (Figure 1B). We tested whether SGO-1 is required for meiotic prophase checkpoints by introducing this allele into *syp-1* mutants (Figure 1A). SYP-1 is required for SC assembly, and in its absence, homologous chromosomes fail to synapse and undergo meiotic recombination [26]. As a result, *syp-1* mutants activate two meiotic checkpoints, the synapsis checkpoint and the DDR, which produces high germline apoptosis (Figure 1C and D) [27]. *syp-1;sgo-1(tm2443)* double mutants had reduced germline apoptosis, suggesting inactivation of either the synapsis checkpoint or the DDR (Figure 1D). Since CEP-1 is required for DDR-induced germline apoptosis [28, 29], elevated apoptosis in *syp-1;cep-1* double mutants is strictly due to synapsis checkpoint activity. To test whether *sgo-1* was a synapsis checkpoint component, we generated *syp-1;cep-1;sgo-1(tm2443)* triple mutants. *syp-1;cep-1;sgo-1(tm2443)* triple mutants had wildtype levels of germline apoptosis when compared to *syp-1;cep-1* double mutants (Figure 1D). This indicates that SGO-1 acts in the synapsis checkpoint and more specifically, the region lost in *sgo-1(tm2443)* mutants is required for its function in the synapsis checkpoint (Figure 1A).

**Figure 1:**
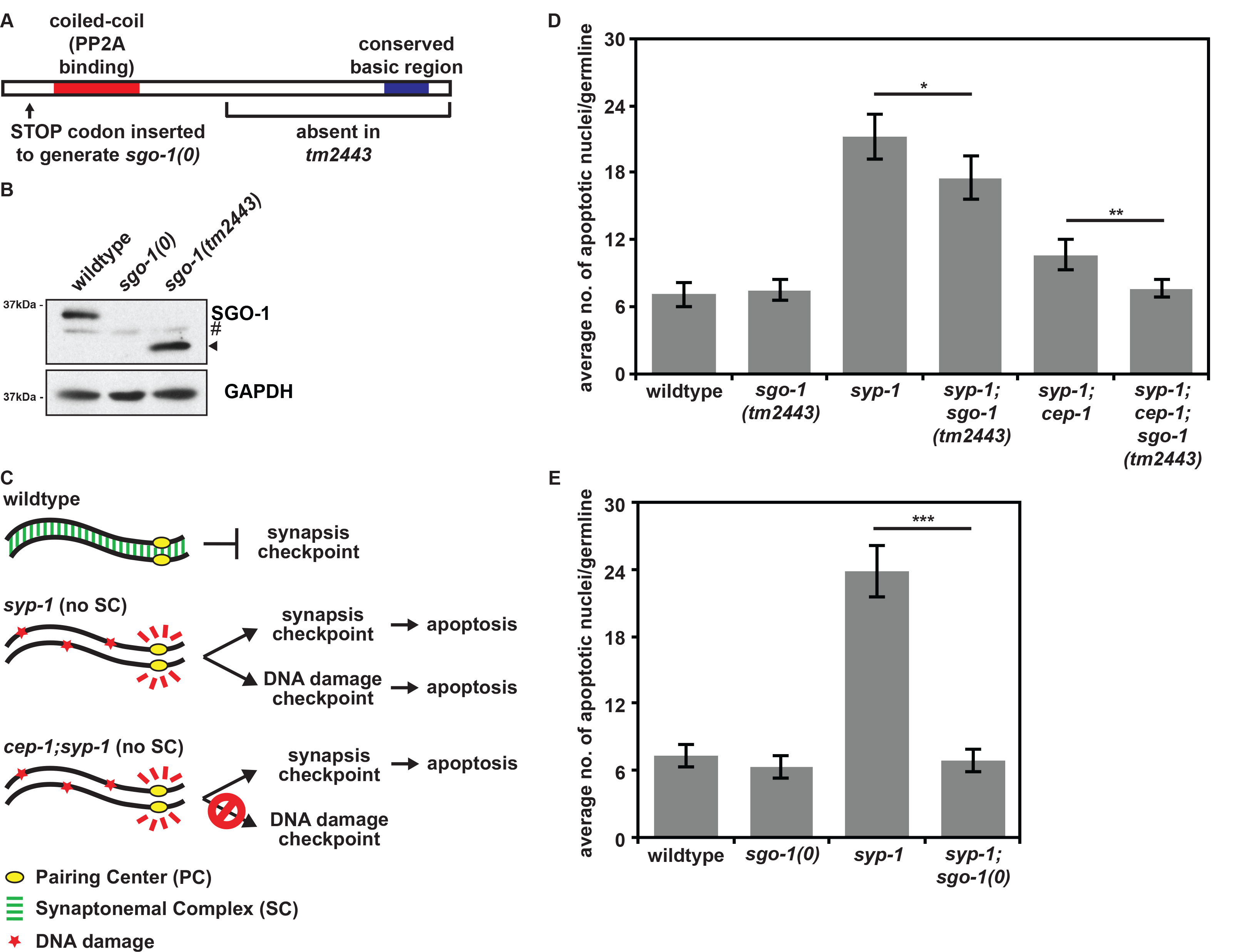
SGO-1 is required for the synapsis checkpoint and the DNA damage response. **A**. Cartoon of SGO-1 protein with relevant mutations indicated. **B**. *sgo-1(0)* mutants have no functional SGO-1 protein expression. Lysates from wildtype, *sgo-1(0)* mutants and *sgo-1(tm2443)* mutants blotted with antibodies against SGO-1 and GAPDH as a loading control. # indicates a background band present in all samples. Arrowhead indicates the truncated version of SGO-1 expressed in *sgo-1(tm2443)* mutants. **C**. Cartoon of meiotic checkpoint activation in *C. elegans*. **D**. A partial loss of function allele of *sgo-1 (sgo-1[tm2443])* reduces apoptosis in *syp-1* single mutants and *syp-1;cep-1* double mutants. **E**. A null mutation in *sgo-1 (sgo-1[0])* reduces apoptosis to wildtype levels in *syp-1* mutants. Error bars indicate 2XSEM. Significance was assessed by performing t-tests. In all graphs, a * indicates a p value < 0.05, a ** indicates a p value < 0.01, and a *** indicates a p value < 0.0001.

To verify this, we introduced the *sgo-1(tm2443)* mutant allele into *meDf2* mutants. *meDf2* is a deletion of the X chromosome Pairing Center (PC) [30], which is required for pairing, synapsis and the synapsis checkpoint [27, 31]. Although *meDf2* homozygotes fail to synapse X chromosomes due to the absence of PCs, they also cannot signal to the synapsis checkpoint, instead activating apoptosis via the DDR [27]. In contrast, the presence of an active PC on unsynapsed X chromosomes in *meDf2* heterozygotes *(meDf2/+)* produces elevated apoptosis via the synapsis checkpoint but not the DDR (Figure S1A) [27]. Consistent with *sgo-1(tm2443)* mutants specifically abolishing the synapsis checkpoint and not the DDR, *meDf2;sgo-1(tm2443)* double mutants had similar levels of apoptosis as *meDf2* single mutants while apoptosis was reduced in *meDf2/+;sgo-1(tm2443)* double mutants in comparison to *meDf2/+* single mutants (Figure S1B).

We wondered if null mutations in *sgo-1* would produce similar results. Therefore, we introduced a stop codon by CRISPR/Cas9 genome editing technology 63 base pairs after the start of the *sgo-1* gene (Figure 1A) and verified that these mutants did not produce SGO-1 protein (Figure 1B). We introduced this null mutation, *sgo-1(0)*, into *syp-1* mutants and were surprised to find that *syp-1;sgo-1(0)* double mutants exhibited wildtype levels of apoptosis (Figure 1E), indicating that SGO-1 function is required for both meiotic checkpoints. Consistent with this analysis, the *sgo-1(0)* mutant allele also reduced apoptosis in both *meDf2* homozygotes and heterozygotes (Figure S1C). Thus, when SGO-1 function is completely abrogated, both the synapsis checkpoint and the DDR are inactive.

### SGO-1 regulates meiotic checkpoint function independent of spindle checkpoint components and PCH-2

We previously identified additional genes that are required for the synapsis checkpoint and showed that they inhibit synapsis in two independent pathways [32]. One pathway involves the microtubule motor, dynein, which is essential for synapsis in *C. elegans* [33]. We previously demonstrated that spindle checkpoint genes, Mad1, Mad2 and Bub3, enforce this requirement for dynein: loss of function mutations in these spindle checkpoint genes restore synapsis when dynein function is knocked down, potentially implicating these factors in a tension-sensing mechanism at PCs [4]. Shugoshin has been shown to respond to changes in tension at centromeres, specifically during biorientation of chromosomes on mitotic or meiotic spindles [34–38]. Further, in humans, mice and Xenopus, Shugoshin interacts directly with Mad2 [39, 40], suggesting that SGO-1 may act with Mad1, Mad2 and Bub3 to regulate the synapsis checkpoint. We tested whether SGO-1 may also be involved in tension-sensing during synapsis by performing RNA interference against the gene that encodes dynein light chain *(dlc-1)* in wildtype, *sgo-1(tm2443), sgo-1(0)* and *mad-1 (mdf-1* in *C. elegans)* null mutants *(mad-1[0])*. To visualize synapsis in these mutants, we performed immunofluorescence against the SC components HTP-3 and SYP-1 (Figure 2A). 76% of germlines from *dlc-1^RNAi^* animals exhibited asynapsis (Figure 2B), visible as meiotic chromosomes with HTP-3 but devoid of SYP-1 (see dashed line in Figure 2A). *mad-1(0);dlc-1^RNAi^* worms suppressed asynapsis in *dlc-1^RNAi^* animals and significantly reduced the percentage of germlines with asynapsis to 24% (Figure 2B) [4]. By contrast and similar to *dlc-1^RNAi^* animals, both *sgo-1(tm2443);dlc-1^RNAi^* and *sgo-1(0);dlc-1^RNAi^* worms (Figure 2A and data not shown) had 75% and 74%, respectively, of germlines with unsynapsed chromosomes (Figure 2B), indicating that SGO-1 does not monitor or regulate meiotic synapsis in the same pathway as Mad1, Mad2 or Bub3.

**Figure 2:**
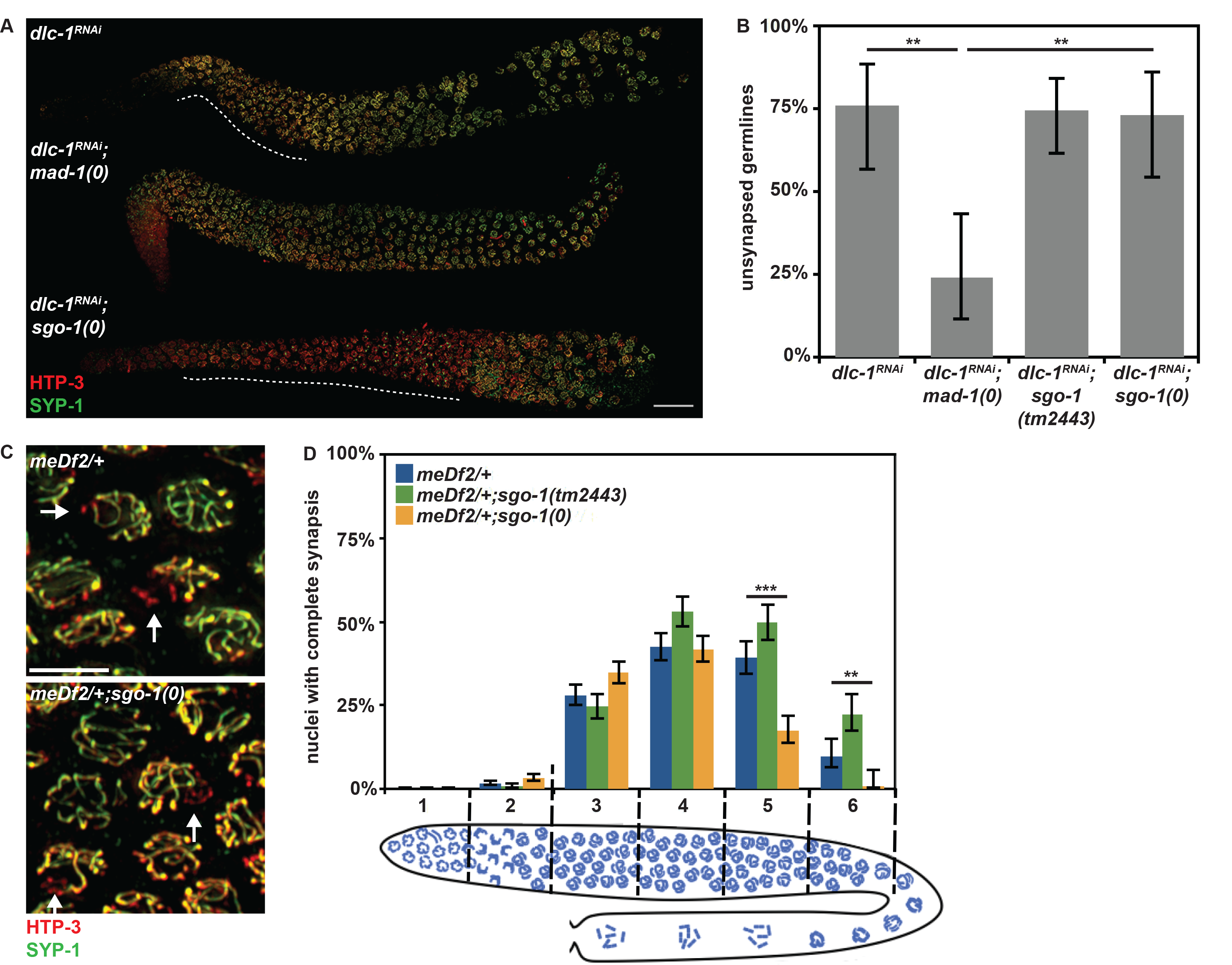
SGO-1 acts in a pathway independent of spindle checkpoint components and PCH-2. **A**. Germlines stained with antibodies against HTP-3 and SYP-1. Dashed white lines highlight regions of asynapsis and scale indicates 20 micrometers. **B**. Mutations in *sgo-1*, unlike loss of *mad-1*, do not suppress asynapsis in *dlc-1^RNAi^* mutants. Error bars indicate 95% confidence intervals. **C**. Meiotic nuclei stained with antibodies against HTP-3 and SYP-1. Unsynapsed chromosomes identified by arrows and scale bar indicates 4 micrometers. **D**. Mutations in *sgo-1* do not suppress asynapsis in *meDf2/+* mutants. In all graphs that include a cartoon depiction of the *C. elegans* germline, meiotic progression is from left to right. Significance was assessed by performing Fisher’s exact test.

The second pathway that inhibits synapsis involves the conserved ATPase, PCH-2 [32]. Similar to mutation of SGO-1, loss of PCH-2 does not suppress the defect in synapsis observed when dynein activity is knocked down [4]. However, loss of PCH-2 rescues the defect in synapsis observed in *meDf2* heterozygotes *(meDf2/+)*, suggesting that PCH-2 inhibits synapsis from non-PC sites [32]. We evaluated synapsis in *meDf2/+, meDf2/+;sgo-1(tm2443)* and *meDf2/+;sgo-1(0)* mutants (Figure 2C and data not shown). We took advantage of the spatio-temporal organization of meiotic nuclei in the germline, dividing the germline into six equivalently sized zones (see cartoon in Figure 2D), and quantified the percentage of nuclei that had competed synapsis. We could not detect any effect on the progression of synapsis in the double mutants (Figure 2D, zones 2 and 3). Instead, we observed what appeared to be more rapid SC disassembly in *meDf2/+;sgo-1(0)* mutants (Figure 2D, zones 5 and 6).

In addition to its effect on synapsis, loss of PCH-2 stabilizes pairing intermediates [32]. We hypothesize that this stabilization of pairing, particularly at PCs, satisfies the synapsis checkpoint in *pch-2;syp-1* double mutants [32]. To assay pairing, we localized the X chromosome PC protein, HIM-8, in *syp-1* mutants (Figure S2A), which allows us to visualize pairing intermediates in the absence of synapsis [26, 41]. We then quantified the percentage of meiotic nuclei that had a single HIM-8 focus, indicating that X chromosomes had paired, as a function of meiotic progression in *syp-1* single mutants as well as *syp-1;sgo-1(tm2443)* and *syp-1;sgo-1(0)* double mutants (Figure S2B). Unlike what we observe in *pch-2;syp1* mutants [32], the progression of pairing in *syp-1;sgo-1(tm2443)* and *syp-1;sgo-1(0)* double mutants was indistinguishable from *syp-1* single mutants (Figure S2B). Altogether, these data suggest that SGO-1 also does not act in the same pathway as PCH-2. Therefore, SGO-1 identifies a third, alternate pathway that regulates synapsis checkpoint function.

### A null mutation in *sgo-1* resembles a partial loss of function allele in the meiotic HORMAD, HTP-3

To determine whether loss of SGO-1 had any effect on synapsis, we monitored synapsis in wildtype worms and *sgo-1* single mutants (Figure 3A) as a function of meiotic progression (Figure 3B), similar to our experiment in Figure 2D. Unlike *mad-1, bub-3* or *pch-2* mutants [4, 32], we did not detect an acceleration of SC assembly (Figure 3B, zones 2 and 3). Instead, similar to what we observed in *meDf2/+sgo-1(0)* double mutants (Figure 2D), we observed that SC disassembly was slightly more rapid in *sgo-1(tm2443)* and significantly more rapid in *sgo-1(0)* mutants than wildtype (see unsynapsed chromosomes in *sgo-1(tm2443)* and *sgo-1(0)* in Figure 3A and Figure 3B, zones 5 and 6).

**Figure 3:**
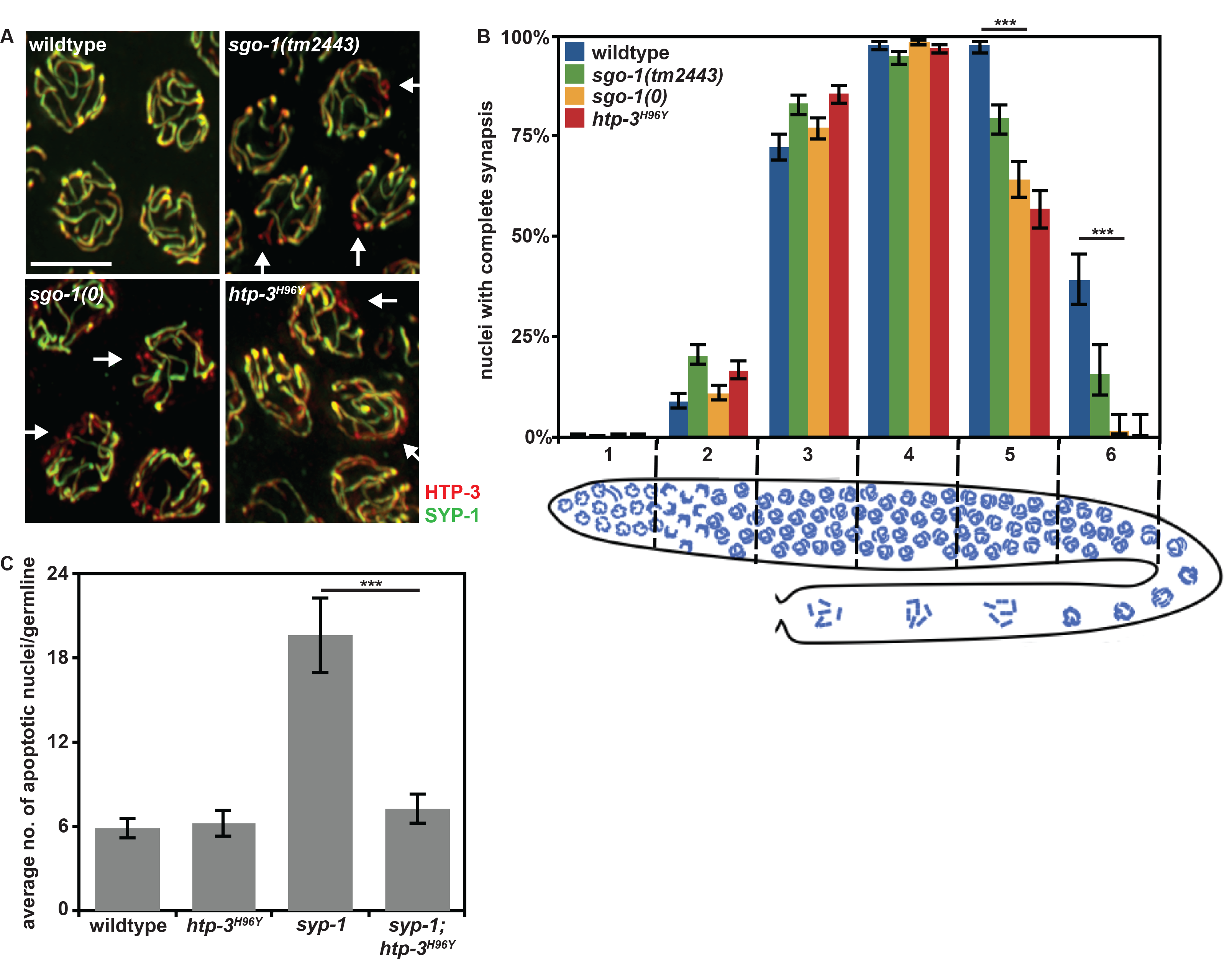
sgo-1 mutants resemble a hypomorphic mutation in htp-3. **A**. Meiotic nuclei in late pachytene stained with antibodies against SC components HTP-3 and SYP-1. Arrows identify chromosomes undergoing SC disassembly. Scale bar indicates 4 micrometers. **B**. Mutations in *sgo-1* accelerate SC disassembly, similar to a partial loss of function mutation in htp-3 *(htp-3^H96Y^)*. Error bars indicate 95% confidence intervals. Significance was assessed by performing Fisher’s exact test. **C**. A partial loss of function mutation in htp-3 *(htp-3^H96Y^)* reduces apoptosis to wildtype levels in *syp-1* mutants. Error bars indicate 2XSEM. Significance was assessed by performing t-tests.

This phenotype reminded us of the reported phenotype of a partial loss of function mutant allele of the meiotic HORMAD, *htp-3, htp-3^H96Y^* (Figures 3A and B) [43]. This mutation converts a histidine at position 96 of the HTP-3 protein to a tyrosine. This amino acid lies in the HORMA domain and is not conserved but resides next to two invariant residues shared between the four meiotic HORMA domain containing proteins in *C. elegans* (HTP-3, HIM-3, HTP-1, and HTP-2), suggesting it might affect HORMA domain function. Given that *sgo-1* mutants resemble *htp-3^H96Y^* mutants in the context of SC disassembly (Figure 3B), and we showed that a subset of meiotic HORMADs are required for checkpoint-induced germline apoptosis [42], we tested what effect this allele had on meiotic checkpoint activation by introducing it into *syp-1* mutants. We found that mutation of the HORMA domain abolished both the synapsis checkpoint and the DDR (Figure 3C), similar to null mutations in *htp-3, him-3* [42] and *sgo-1* (Figure 1D). Thus, both *htp-3^H96Y^* and *sgo-1(0)* mutants abrogate meiotic checkpoint function and prematurely disassemble the SC, suggesting they act in the same pathway.

### SGO-1 limits non-homologous DNA repair and promotes crossover assurance

HTP-3^H96Y^ also affects the progression of DNA repair [43] by inappropriately activating non-homologous DNA repair mechanisms. We tested the role of SGO-1 in meiotic recombination. We focused these experiments on the null mutation of *sgo-1*, since this allele also affected the DDR and exhibited additional phenotypes that most closely resembled *htp-3^H96Y^* mutants (Figures 1E and 3).

First, we monitored the progression of DNA repair. For this experiment, we performed immunofluorescence against the DNA repair factor RAD-51 (Figure 4A). RAD-51’s appearance on meiotic chromosomes indicates the formation of double strand breaks and its disappearance shows entry into a DNA repair pathway [44]. When we follow the dynamics of RAD-51 appearance and disappearance in wildtype and *sgo-1(0)* single mutants, they are exceedingly similar (Figure 4B). However, when we did this experiment in the *syp-1* mutant background, in which the inability to synapse prevents DNA repair from using a homologous chromosome as a template [44], we saw that *syp-1;sgo-1(0)* double mutants had sharply reduced average number of RAD-51 foci, particularly in zone 5, suggesting these double mutants repair double strand breaks more rapidly than *syp-1* single mutants (Figure 4B).

**Figure 4:**
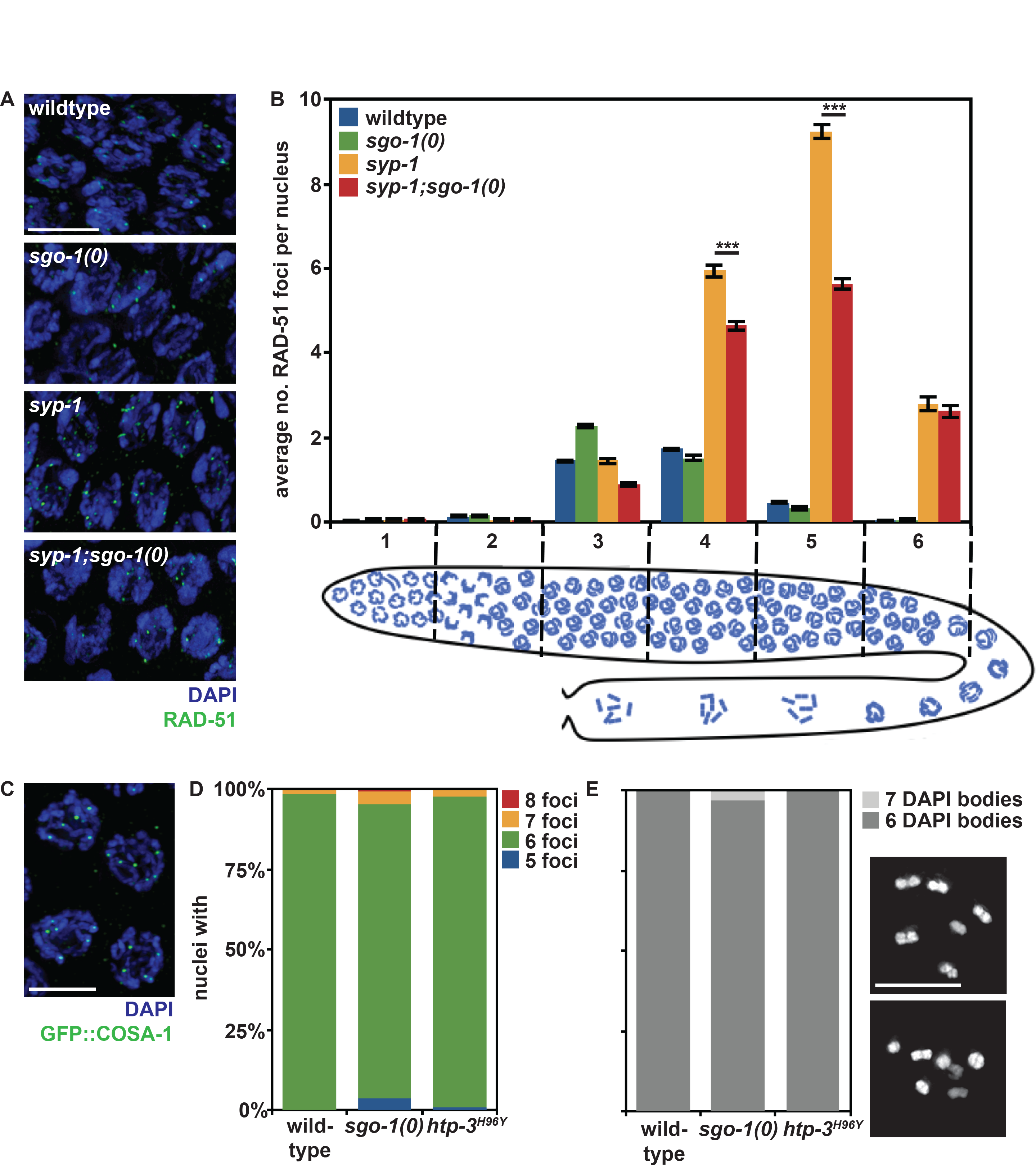
SGO-1 is not essential for crossover formation but is required to promote interhomolog DNA repair and crossover assurance. **A**. Meiotic nuclei stained with DAPI and antibodies against RAD-51. Error bars indicate 2XSEM. **B**. Homolog-independent mechanisms of DNA repair are active in *syp-1;sgo-1(0)* double mutants. In all images, scale bar indicates 4 micrometers. Significance was assessed by performing t-tests. **C**. Meiotic nuclei stained to visualize DNA (DAPI) and GFP::COSA-1. **D**. The percentage of meiotic nuclei with 5, 6, 7 and 8 GFP::COSA-1 foci. **E**. The percentage of meiotic nuclei with 6 (top image) or 7 DAPI staining bodies (bottom image).

DSB-1 and DSB-2 localize to chromosomes and are required for the formation of double strand breaks. When meiosis is defective, DSB-1 and DSB-2 remain on chromosomes [45, 46] and their persistence depends on the recruitment of a subset of meiotic HORMADs to chromosomes [5]. To determine whether the reduction in RAD-51 foci in *syp-1;sgo-1(0)* double mutants is because *sgo-1(0)* mutants fail to activate this meiotic feedback mechanism, we visualized DSB-1 and DSB-2 in *syp-1* and *syp-1;sgo-1(0)* double mutants and could detect no difference in their staining pattern (data not shown). From these data, we conclude that SGO-1 typically prevents the activation of alternate DNA repair mechanisms, such as using the sister chromatid as a repair template, to promote homologous DNA repair.

We reasoned that this effect on DNA repair might have consequences on crossover formation. We monitored crossover formation by evaluating both GFP::COSA-1 localization and bivalent formation. COSA-1 localizes to presumptive crossovers in late meiotic prophase (Figure 4C). The six pairs of chromosomes in *C. elegans* exhibit crossover assurance, in which every pair of chromosomes has at least one crossover, and strict crossover control, in which every pair of chromosomes enjoys only a single crossover. As a result, we observe 6 GFP::COSA-1 foci in greater than 98% of meiotic nuclei in wildtype animals (Figure 4B). *sgo-1(0)* mutants show a significant increase (p value < 0.01, Fisher’s exact test) in nuclei with five GFP::COSA-1 foci, indicating a subtle loss of crossover assurance (Figure 4D). Interestingly, we could not detect a loss of crossover assurance in *htp-3^H96Y^* mutants (Figure 4D), suggesting that either the requirement for SGO-1 in regulating meiotic DNA repair might be stronger than that of a single meiotic HORMAD or HTP-3^H96Y^ might still retain some activity.

Next, we assessed bivalent formation. In wildtype nuclei and *htp-3^H96Y^* mutants, all chromosome pairs are linked by chiasmata in late meiotic prophase and we always see 6 DAPI stained bodies (Figure 4E, top image). We observed non-recombinant chromosome pairs, or univalents (Figure 4E, bottom image), in *sgo-1(0)* mutants in 3% of meiotic nuclei in late prophase (p value < 0.05, Fisher’s exact test), verifying the subtle loss of crossover assurance in this mutant background (Figures 4D and SE).

### SGO-1 promotes the recruitment of HUS-1::GFP to sites of DNA damage

Given the effect that loss of SGO-1 has on meiotic DNA repair and recombination, we wondered if SGO-1’s role in the DDR could be involved in recruiting early DDR components. An early event in DDR is the recruitment of the conserved 9-1-1 complex, which includes the factors MRT-2 (the *C. elegans* Rad1 ortholog), HPR-9 (the *C. elegans* Rad9 ortholog) and HUS-1, to sites of damage [47, 48]. To visualize recruitment of the 9-1-1 complex, we localized HUS-1::GFP in wildtype, *sgo-1(0), htp-3^H96Y^, syp-1, syp-1;sgo-1(0)* and *syp-1;htp-3^H96Y^* mutants (Figure 5A). Wildtype meiotic nuclei had very few HUS-1::GFP foci (Figure 5B). Both *sgo-1(0)* and *htp-3^H96Y^* single mutants exhibited slightly more HUS-1::GFP foci (Figure 5B), indicating that the DDR is weakly active in these backgrounds despite normal levels of apoptosis (Figures 1E and 3C). This may reflect the inappropriate activation of non-homologous DNA repair in these mutants, despite the apparent normal progression of DNA repair ([43] and Figure 4E). Meiotic nuclei in *syp-1* single mutants displayed many more HUS-1::GFP foci (Figure 5B). By contrast, we observed a sharp reduction in the average number of HUS-1::GFP foci in *syp-1;sgo-1(0)* and *syp-1;htp-3^H96Y^* double mutants (Figure 5B), albeit not to the average numbers we observed in the single mutant backgrounds. This defect in the ability to recruit HUS-1::GFP is entirely consistent with the reduction in DDR-induced apoptosis we also detected in these double mutants (Figures 1E, S1C and 3C). Thus, SGO-1 is required to robustly recruit components of the 9-1-1 complex, acting early in the meiotic DDR.

**Figure 5:**
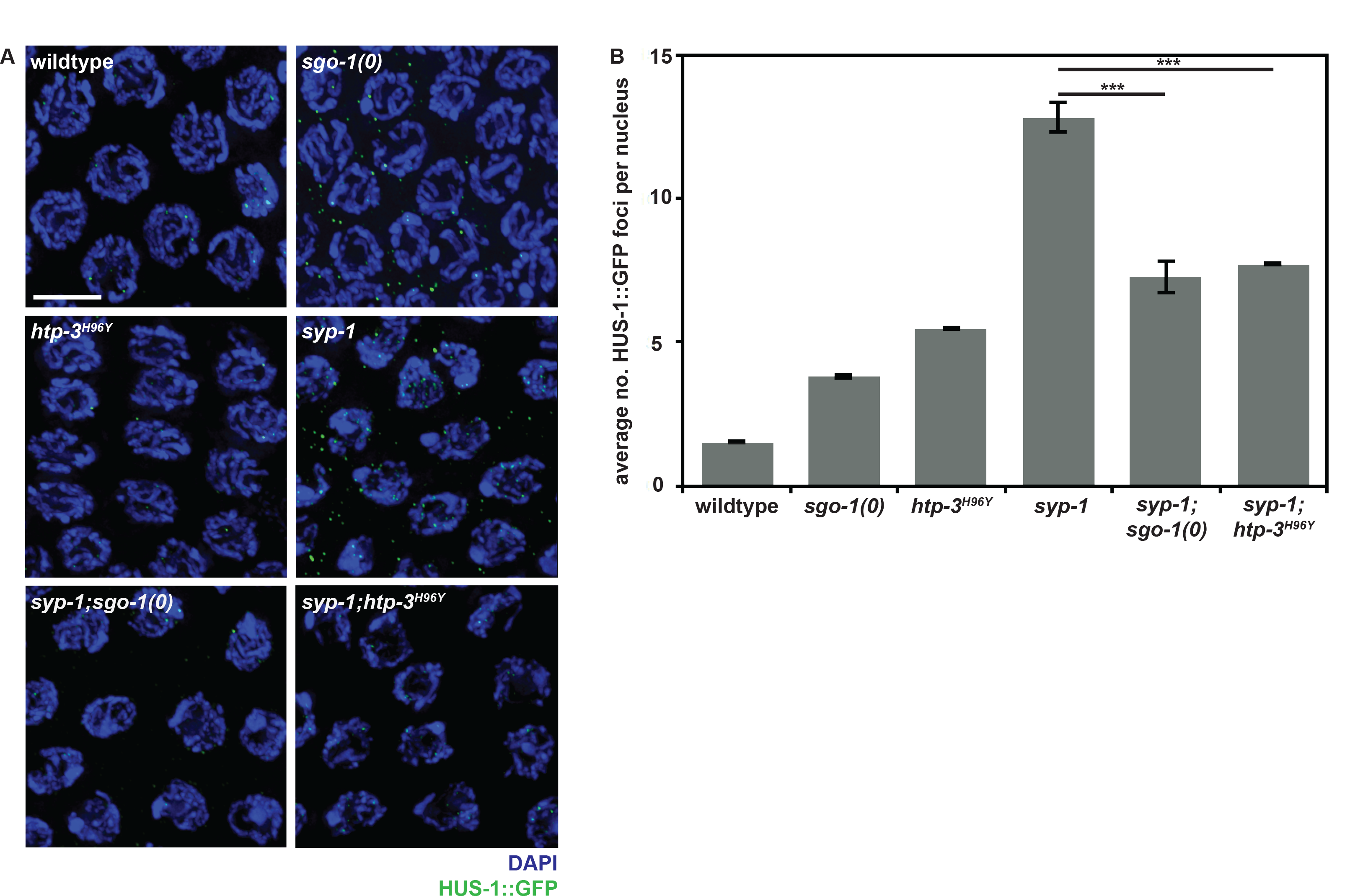
SGO-1 is required for robust localization of HUS-1::GFP to sites of DNA damage. A. Meiotic nuclei stained to visualize DNA (DAPI) and HUS-1::GFP. Scale bar indicates 5 micrometers. **B**. SGO-1 and HTP-3 function are required for robust recruitment of HUS-1::GFP when the DNA damage checkpoint is active. Error bars indicate 2XSEM. Significance was assessed by performing t-tests.

### SGO-1 localizes to premeiotic and late meiotic prophase nuclei

We localized the SGO-1 protein in the hermaphrodite germline. To our surprise its staining was limited to nuclei just prior to entry into meiotic prophase, which are often defined as pre-meiotic, and in late meiotic prophase (Figure 6A). HTP-3 was also present in these pre-meiotic nuclei but was not yet visibly assembled into chromosome axes, suggesting that SGO-1 may be regulating early events in axis morphogenesis. Upon the appearance of discrete HTP-3 axes in early prophase nuclei, SGO-1 protein was conspicuously absent (Figure 6B). When the SC undergoes ordered disassembly in diplotene of meiotic prophase, SGO-1 reappears in meiotic nuclei (Figure 6C). This localization pattern was unchanged in *sgo-1(tm2443)* mutants and absent in *sgo-1(0)* mutants (data not shown).

**Figure 6:**
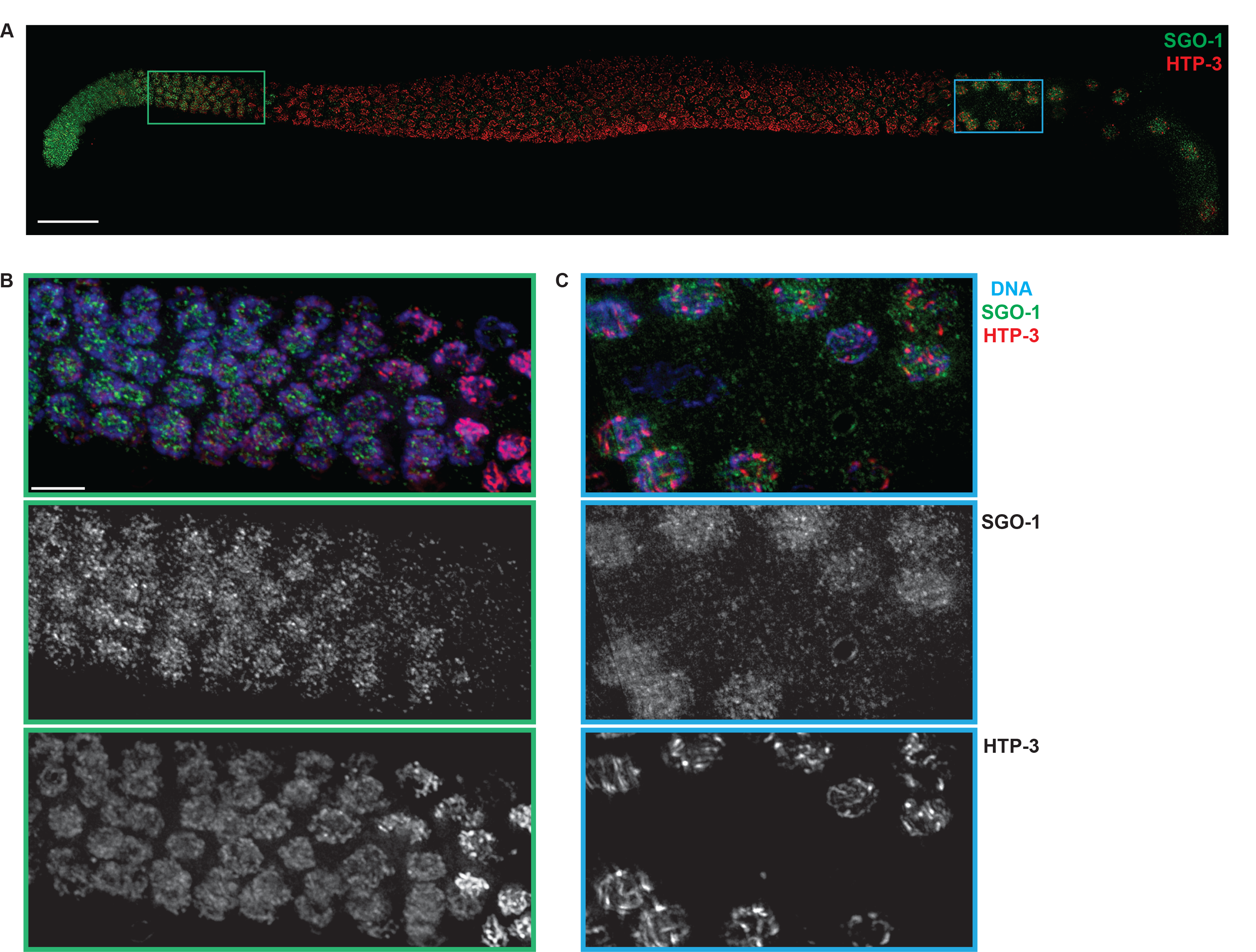
SGO-1 localizes to pre-meiotic nuclei. **A**. Wildtype germline stained with antibodies against SGO-1 and HTP-3. Scale bar indicates 20 micrometers. Pre-meiotic nuclei (**B**) and nuclei in late meiotic prophase (**C**) stained with DAPI and antibodies against SGO-1 and HTP-3. Scale bars in B and C indicates 4 micrometers.

## Discussion

The phenotypes we have characterized when Shugoshin function is compromised or abolished in *C. elegans* are highly reminiscent of mutations in cohesin or other meiotic axis components, namely the inability to activate meiotic checkpoints [4–10], the loss of crossover assurance, and the activation of homolog-independent, presumably sister chromatid-dependent, DNA repair mechanisms (reviewed in [3]). Indeed, we show that *sgo-1* null mutants resemble a loss of function mutation in the meiotic axis component and HORMAD protein, HTP-3 [43]. Despite being dispensable for normal pairing and synapsis in *C. elegans* (Figures S2 and 3B), we propose that SGO-1 is required to generate meiotic chromosome architecture competent for checkpoint activation and the normal progression of meiotic recombination (Figure 7). Further, we hypothesize that this role is conserved but unappreciated given the focus on Shugoshin’s role in regulating two-step loss of sister chromatid cohesion during meiotic chromosome segregation. The requirement for Shugoshin in maintaining meiotic synapsis in rice, a phenotype startlingly similar to the premature SC disassembly we detect in *sgo-1* mutants, strongly supports this possibility [49]. More importantly, our findings expand the repertoire of Shugoshin’s functions in controlling chromosome segregation beyond being a platform or adapter protein at centromeric regions.

**Figure 7:**
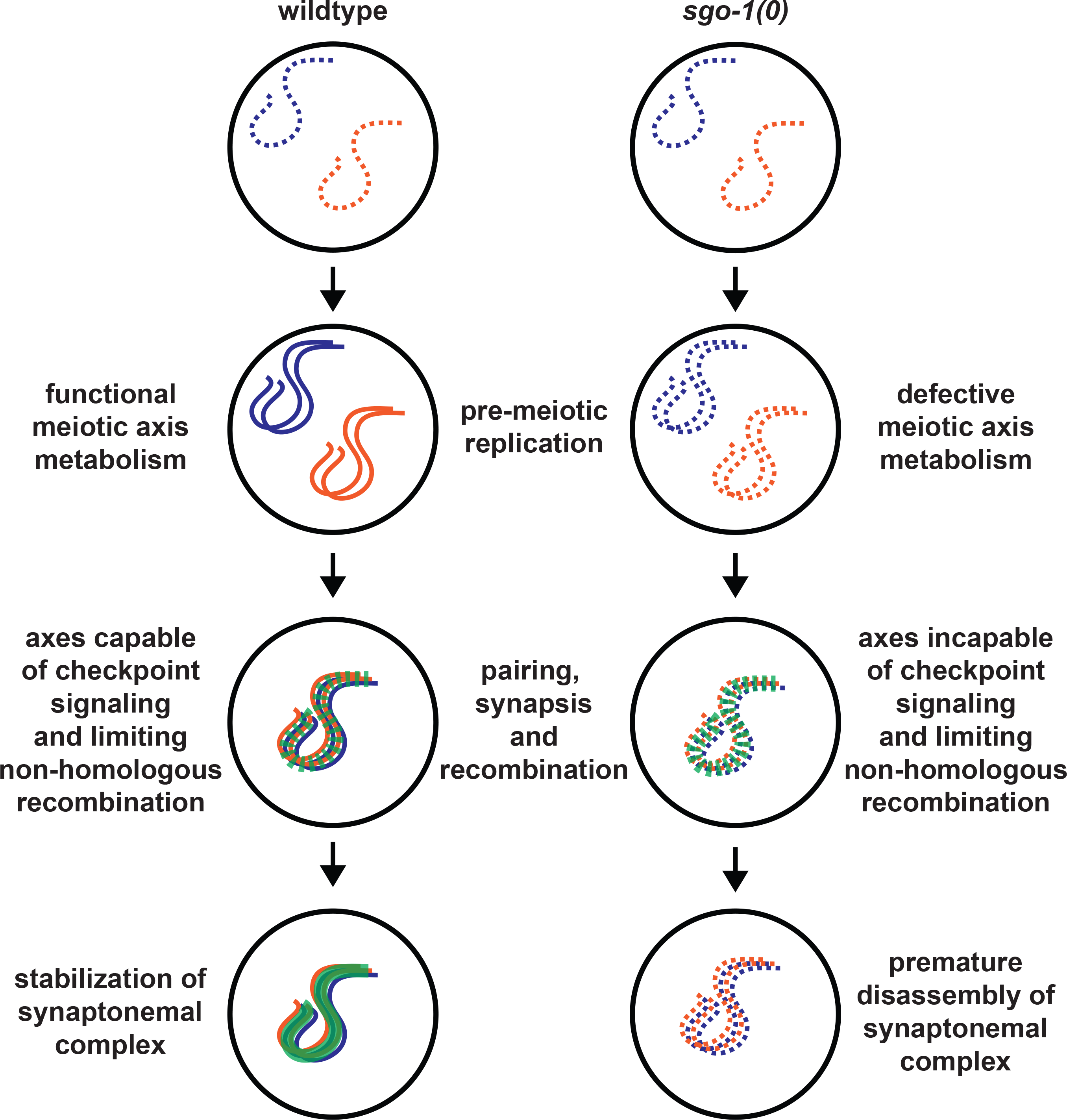
Model for SGO-1 function in meiotic prophase. Orange and blue dotted lines indicate homologous chromosomes prior to replication. During pre-meiotic replication in wildtype nuclei, meiotic axis metabolism initiates (continuous blue and orange lines), generating axes functional for pairing, synapsis (dotted green line), checkpoint signaling, limiting non-homologous recombination and eventual stabilization of the SC (solid green line). *sgo-1* mutants fail to form fully functional axes (represented by the continued presence of dotted blue and orange lines), resulting in a loss of checkpoint signaling, the activation of non-homologous recombination mechanisms and premature disassembly of the SC.

Given our proposal that SGO-1 acts in the same pathway as meiotic HORMADs for meiotic checkpoint function, we were surprised to see that *sgo-1* mutants did not resemble *pch-2* mutants (Figures 2C, 2D, S2 and 3B). In budding yeast and mice, PCH-2, and its mammalian ortholog TRIP13, regulate meiotic HORMADs in a feedback mechanism that signals proper meiotic progression [7, 10, 14, 50]. This discrepancy may be because meiotic HORMADs regulate meiotic checkpoint function through multiple mechanisms, one involving PCH-2 and one involving SGO-1. Further, our identification of at least three separate pathways that trigger germline apoptosis in response to defects in synapsis [4, 42, this study] demonstrates the stringency of the synapsis checkpoint in *C. elegans*, presenting a marked contrast to oogenesis in mammalian cells [51]. This stringency may reflect the importance of synapsis to the generation of chiasma [44] or the regulation of germline apoptosis in *C. elegans*.

The localization of SGO-1 in the pre-meiotic zone corresponds to when the meiosis specific cohesin complex defined by the kleisin, REC-8, is loaded onto meiotic chromosomes [52] and pre-meiotic replication occurs. However, it precedes when the meiosis specific cohesin complex defined by the kleisins, COH-3/COH-4, is loaded onto meiotic chromosomes [53]. Therefore, we suggest that SGO-1’s role in promoting fully functional chromosome axes is likely associated with pre-meiotic replication and involves cohesin, specifically cohesin complexes defined by REC-8. In addition to SGO-1’s localization, this is based on its characterized role regulating sister chromatid cohesion during meiotic chromosome segregation [19–21], the reported biochemical interaction between meiotic HORMADs and cohesin [11, 54] and the observation that complete loss of cohesin function also affects the ability to recruit HUS-1::GFP [6]. Unfortunately, we could not detect any obvious defects in the localization of REC-8, COH-3 or COH-4 in *sgo-1(0)* mutants (data not shown). Shugoshin also regulates additional factors required for chromosome structure and function, such as condensin [55, 56], raising the possibility that Shugoshin’s effect on meiotic prophase events occurs through factors in addition to or independent of cohesin.

SGO-1’s localization is similar to that of the cohesin regulator WAPL-1, whose early localization during pre-meiotic replication also affects meiotic axis structure [57]. Based on this colocalization and reports that Shugoshin and Wapl may antagonize each other [58], we tested whether we could detect a genetic interaction between mutations in *wapl-1* and *sgo-1*, specifically whether loss of *sgo-1* would suppress the reduction in meiotic axis length observed in *wapl-1* mutants. However, meiotic chromosomes in *wapl-1;sgo-1* double mutants resembled those in *wapl-1* single mutants (data not shown), suggesting that these two factors appear not to antagonize each other when regulating meiotic axis structure.

The *tm2443* mutant allele acts as a separation of function allele (Figures 1 and S1), indicating that SGO-1’s role in the synapsis checkpoint and the DDR are separable. Two functional portions of SGO-1 are absent in the protein produced by the *tm2443* allele and are potentially essential for the synapsis checkpoint but dispensable for the DDR. These include the highly conserved basic “SGO motif,” which mediates binding to histone H2A phosphorylated by the conserved cell cycle kinase and spindle checkpoint component Bub1 [59]. Given our interest in roles for spindle checkpoint components in regulating and monitoring meiotic synapsis [4], testing whether Bub1 and its kinase activity are required for the synapsis checkpoint is a current focus. The middle section of the protein that is highly divergent in both length and sequence is also missing in the protein generated by the *tm2443* mutant allele. This region appears to mediate interactions with a wide array of proteins, including cohesin, Mad2 and the microtubule motor, MCAK [39, 40, 58, 60]. Considering that loss of *sgo-1* fails to suppress the synapsis defect when the microtubule motor dynein is knocked down (Figures 2A and B), unlike loss of Mad2 [4], and that SGO-1 is enriched in the nucleus (Figure 6), where microtubules are not present [33], we think it unlikely that either interaction with Mad2 or MCAK explains SGO-1’s function in the synapsis checkpoint. This region is also subject to phospho-regulation by important cell cycle kinases in some organisms (reviewed in [61]), raising the possibility that regulation of this portion of Shugoshin contributes to its synapsis checkpoint role. The N-terminal coiled-coil region, which has been shown to promote dimerization [62] as well as interact with both PP2A phosphatase [63] and the chromosome passenger complex (CPC) [64], may be required for Shugoshin’s role in the DDR as well as in regulating meiotic recombination. However, the conserved kinase Aurora B, a component of the CPC, is prevented from interacting with meiotic chromosomes in *C. elegans* to promote sister chromatid cohesion [65] and is not required for either the synapsis checkpoint or the DDR (data not shown), ruling out that an interaction with this complex explains SGO-1’s contribution to meiotic axis function. No role for PP2A in meiotic prophase has been reported but it’s possible a role early in meiosis has been overlooked, similar to Shugoshin’s. Additional domain analysis and identification of Shugoshin’s meiotic interactors in *C. elegans* will determine how Shugoshin manages its multiple roles during meiotic prophase.

The premature disassembly of the SC in both *sgo-1* null mutants and *htp-3^H96Y^* mutants provides a potential opportunity to reconcile what previously appeared to be disparate observations in multiple meiotic systems. The stability of axis and SC components on meiotic chromosomes is tightly controlled and linked to the progression of meiotic recombination. In budding yeast and mice, this includes the Pch2/Trip13-dependent redistribution or removal of meiotic HORMADs from chromosomes as chromosomes synapse [14, 50, 66]. Given the multiple roles meiotic HORMADs play during prophase, this redistribution or removal likely accomplishes three things: 1) it prevents additional double strand breaks [7, 67, 68]; 2) it allows any remaining double strand breaks to be repaired using the sister chromatid as a template or by mitotic-like mechanisms [69–72]; and 3) it signals the proper progression of meiotic prophase [7–10]. In budding yeast, central element components of the SC also undergo turnover, but it is limited to regions associated with meiotic recombination [73].

In *C. elegans*, relocalization or redistribution of meiotic HORMADs does not occur until SC disassembly. However, several reports have highlighted how the SC becomes more stable later in meiotic prophase before undergoing ordered disassembly (see Figure 7) [74–77]. This stability relies on the presence of a crossover-specific intermediate in cis [74, 76]: chromosomes that fail to undergo crossover recombination disassemble their SCs prematurely, similar to *sgo-1* null and *htp-3^H96Y^* mutants (Figures 3A, B and [43]). Since this portion of meiotic prophase coincides with a loss of homolog access during DNA repair [78] and a release from meiosis-specific DNA repair mechanisms [79], it seems likely that some modification of axis components also occurs during this period of meiotic prophase. We suggest that this modification may contribute to SC stabilization, and its eventual ordered disassembly, potentially analogous to the remodeling of meiotic HORMADs in budding yeast and mice. SC disassembly is accelerated in *sgo-1* null and *htp-3^H96Y^* mutants despite the presence of crossover-specific recombination intermediates (Figures 3A, B and [43]), suggesting that a fully functional meiotic axis is important for this stabilization (Figure 7). SC disassembly is delayed in *C. elegans pch-2* mutants, implicating this factor in the process, analogous to yeast and mammals [32]. We speculate that this remodeling manifests itself differently in *C. elegans* than in yeast or mice because *C. elegans* relies on synapsis for early events in meiotic recombination, such as ZHP-3 recruitment [80], and uses axis components, including meiotic HORMADs, to direct the two step loss of sister chromatid cohesion [22, 23].

## Materials and Methods

### Genetics and worm strains

The wildtype *C. elegans* strain background was Bristol N2 [81]. All experiments were performed on adult hermaphrodites at 20°C under standard conditions unless otherwise stated. Mutations and rearrangements used were as follows:

LG I: *mnDp66, cep-1(gk138), htp-3(vc75), hus-1(op241)*

LG II: *meIs8 [Ppie-1::GFP::cosa-1 + unc-119(+)]*

LG IV: *sgo-1(tm2443), sgo-1(blt2), nT1[unc-?(n754) let-?(m435)] (IV, V), nTI [qIs51]*

LG V: *syp-1(me17), mad-1(gk2), dpy-11(e224), bcIs39 [lim-7p::ced-1::GFP + lin-15(+)]*

LG X: *meDf2*

*opIs34 [Phus-1::hus-1::GFP + unc-119(+)]*

*meDf2* is a terminal deficiency of the left end of the X chromosome that removes the X chromosome PC as well as numerous essential genes [30]. For this reason, homo- and hemizygous *meDf2* animals also carry a duplication *(mnDp66)* that includes these essential genes but does not interfere with normal X chromosome segregation [82] or synapsis checkpoint signaling [27]. For clarity, it has been omitted from the text.

The *sgo-1* null allele *(sgo-1[0]), blt2*, was created by CRISPR-mediated genomic editing as described in [83, 84]. pDD162 was mutagenized using Q5 mutagenesis and oligos TAAAACTGCAGCATGTGCCGTTTTAGAGCTAGAAATAGCAAGT and CAAGACATCTCGCAATAGG. The resulting plasmid was sequenced and three different correct clones (50ng/ul total) were mixed with pRF4 (120ng/ul) and the repair oligo ATTTGTATTTTACACATAAACTTTGTAAATATAATAATACCTTCTTTAGAGCTAGCTTGGTCGTTTTTTTGCTGCTACAATTCCTCCAAAAATAGATTGTGCAGTTT (30ng/ul). Wildtype worms were picked as L4s, allowed to age 15-20 hours at 20°C and injected with the described mix. Worms that produced rolling progeny were identified and F1 rollers, as well as their wildtype siblings, were placed on plates seeded with OP50, 1-2 rollers per plate and 6-8 non-rolling siblings per plate, and allowed to produce progeny. PCR and NheI digestions were performed on these F1s to identify worms that contained the mutant allele and individual F2s were picked to identify mutant homozygotes. Multiple homozygotes carrying the *sgo-1(blt2)* mutant allele were backcrossed against wildtype worms at least three times and analyzed to determine whether they produced the same mutant phenotype.

### Quantification of Germline Apoptosis

Scoring of germline apoptosis was performed as previously descried in [27] with the following exceptions. L4 hermaphrodites were allowed to age for 22 hours. They were then mounted under coverslips on 1.5% agarose pads containing 0.2mM levamisole for wildtype moving strains or 0.1mM levamisole for *dpy-11* strains. A minimum of twenty-five germlines were analyzed for each genotype.

### Antibodies, Immunostaining and Microscopy

DAPI staining and immunostaining was performed as in [27] 20 to 24 hours post L4 stage. Primary antibodies were as follows (dilutions are indicated in parentheses): rabbit anti-SYP-1 (1:500) [26], chicken anti-HTP-3 (1:1000) [31], guinea pig anti-HIM-8 (1:250)[41], rabbit anti-SGO-1 (1:30,000) [25], mouse anti-GFP (1:100) (Invitrogen) and rabbit anti-RAD-51 (1:5000) [85]. Secondary antibodies were Cy3 anti-rabbit, anti-guinea pig and anti-chicken (Jackson Immunochemicals) and Alexa-Fluor 488 anti-guinea pig and anti-rabbit (Invitrogen). All secondary antibodies were used at a dilution of 1:500. DAPI staining of meiotic nuclei in late meiotic prophase to visualize bivalents was performed 48 hours post-L4 stage.

Quantification of synapsis and RAD-51 foci was performed with a minimum of three whole germlines per genotype as in [41] on animals 24 hours post L4 stage.

All images were acquired using a DeltaVision Personal DV system (Applied Precision) equipped with a 100X N.A. 1.40 oil-immersion objective (Olympus), resulting in an effective XY pixel spacing of 0.064 or 0.040 µm. Three-dimensional image stacks were collected at 0.2-µm Z-spacing and processed by constrained, iterative deconvolution. Image scaling and analysis were performed using functions in the softWoRx software package. Projections were calculated by a maximum intensity algorithm. Composite images were assembled and some false coloring was performed with Adobe Photoshop.

### Westerns

For immunoblotting, samples were run on SDS-PAGE gels, transferred to nitrocellulose, blocked in a PBST + 5% (w/v) non-fat milk solution, and then probed with rabbit anti-SGO-1 (dilution 1:30,000) and anti-GAPDH (1:5000) overnight at 4°C. Blots were washed 3x for 10 minutes in PBST, probed for 1 hour using an HRP-conjugated secondary antibody (rabbit or mouse; GE Healthcare), washed 3x for 10 minutes in PBST, and then analyzed using a chemiluminescent substrate (Thermo Scientific).

### Feeding RNAi

For RNAi *dlc-1^RNAi^* and empty vector (L4440) clones from the Ahringer laboratory [86] were used. Bacteria strains containing *dlc-1^RNAi^* and empty vector controls were cultured overnight in 10ml LB + 50ug/ul carbenicillin, centrifuged, and resuspended in 0.5 ml LB + 50ug/ul carbenicillin. Sixty microliters of the RNAi bacteria was spotted onto NGM plates containing 1mM IPTG + 50ug/ul carbenicillin and allowed to grow at room temperature overnight. L4 hermaphrodite worms were picked into M9, transferred to these plates, allowed to incubate for 2-3 hours and then transferred to fresh RNAi plates to be dissected 48 hours post L4. A minimum of 28 germlines were scored for each genotype.

---

**Figure S1.**
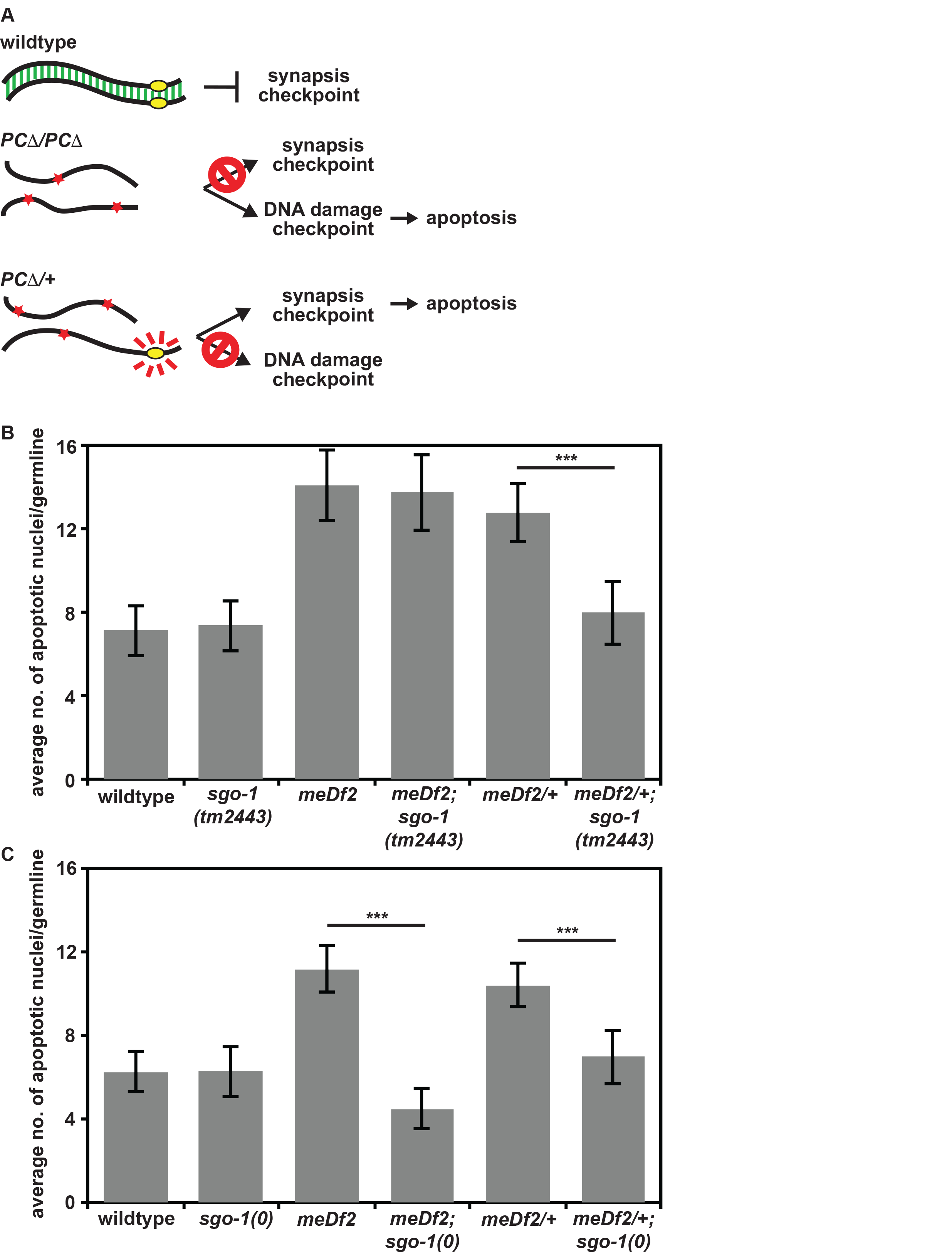
SGO-1 is required for the synapsis checkpoint and the DNA damage response. **A**. Cartoon of meiotic checkpoint activation in *C. elegans*. **B**. A partial loss of function allele of *sgo-1 (sgo-1[tm2443])* reduces apoptosis in *meDf2* heterozygotes *(PCΔ/+)* but not *meDf2* homozygote mutants *(PCΔ/PCΔ)*. **C**. A null mutation in *sgo-1 (sgo-1[0])* reduces apoptosis in both *meDf2* homozygotes *(PCΔ/PCΔ)* and *meDf2* heterozygotes *(PCΔ/+)*. Error bars indicate 2XSEM. Significance was assessed by performing t-tests.

**Figure S2:**
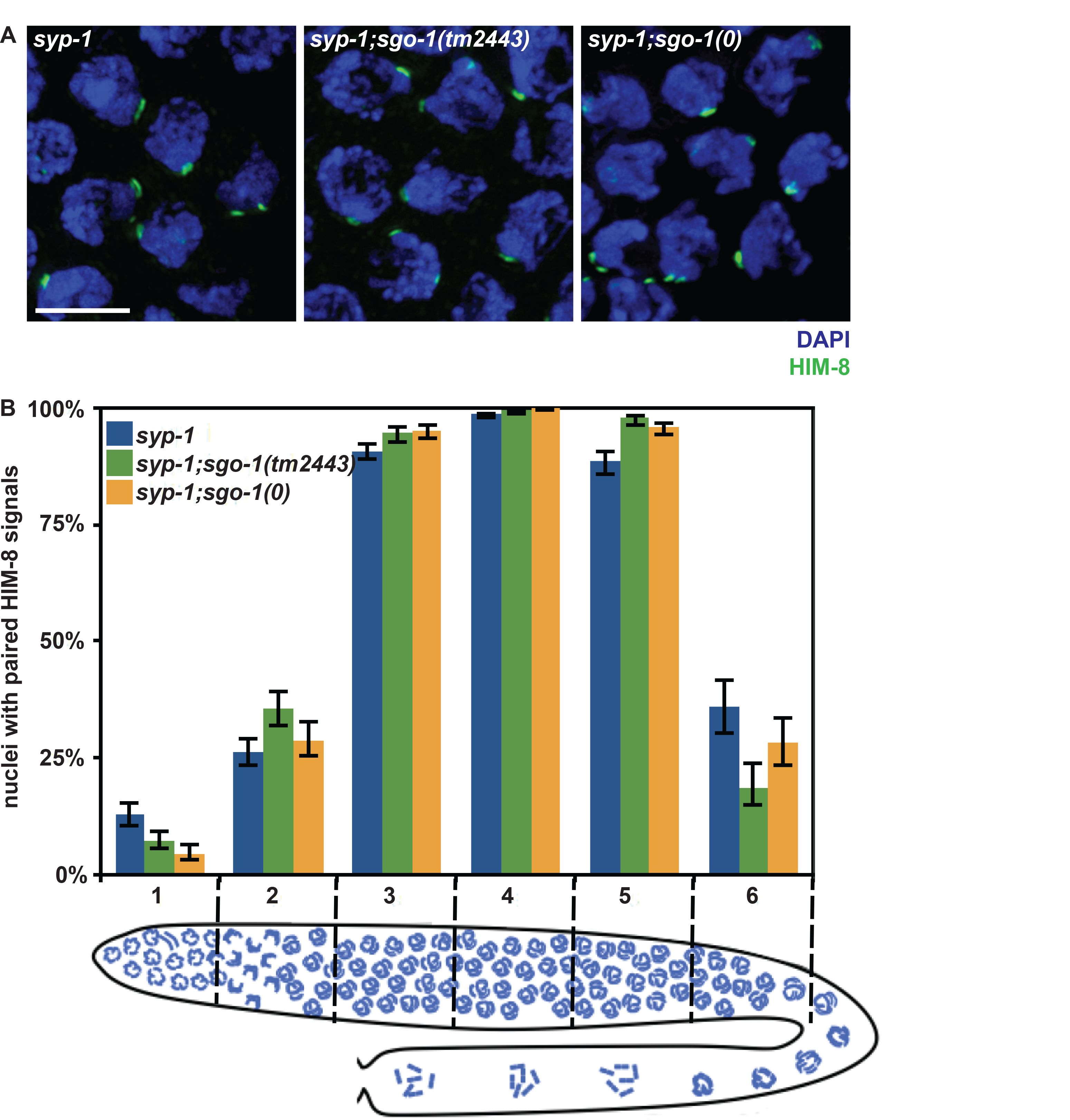
SGO-1 does not regulate pairing. **A**. Meiotic nuclei stained with DAPI and antibodies against HIM-8. Scale bar indicates 4 micrometers. **B**. Pairing in *syp-1;sgo-1* double mutants is indistinguishable from pairing in *syp-1* single mutants. Error bars indicate 95% confidence intervals.

